# The gut microbiome in early age-related macular degeneration

**DOI:** 10.1101/2025.03.25.645132

**Authors:** Beatrix Feigl, Prakash Adhikari, Flavia Huygens, Nadeesha Jayasundara, Andrew J Zele

## Abstract

The gut microbiome is implicated in the development of advanced age-related macular degeneration (AMD) but no study has investigated microbial composition in early AMD nor controlled for important microbiome modulators such as diet and light. This is crucial as diet can change the microbiome rapidly as well as affects disease progression. In addition, the light signalling photoreceptors are dysfunctional in AMD. Here we determined the gut microbiota by conducting 16S DNA metagenomic sequencing of 40 faecal samples from 20 participants with and without early AMD. We normalised both groups to the same diet over 5 days and determined gut microbial composition before and after the diet. To control for light, we assessed habitual light exposure with actigraphy and the post-illumination pupil light response (PIPR) quantified photoreceptor signalling. *Clostridium termitidis, Bacteroides gallinarum*, and *Bacteroides finegoldii* were significantly abundant in early AMD compared to controls at both time points. Interestingly, pro-inflammatory species such as *Sutterella* were only significantly abundant in AMD before the diet. After the diet, the gut microbiome shifted to a significantly greater abundance of commensal bacteria in AMD. Controls and AMD patients had a similar ambient light exposure, however the PIPR was significantly reduced in AMD suggesting impaired light signalling. We are the first to show a distinct gut microbiome in early AMD and infer that dietary changes may positively affect the gut microbiome at early stage of the disease.

## Introduction

In age-related macular degeneration (AMD), the gut microbiome has been implicated of either being protective against or contributing to its progression (Rowan et al. 2017, Andriessen et al. 2016, Zinkernagel et al. 2017). This is thought to be due to a dysregulated inflammatory immune response in the gut leading to dysbiosis (Chen and Xu 2015) (Napolitano et al. 2021, Rinninella et al. 2018). Currently only studies in mice (Rowan et al. 2017, Andriessen et al. 2016) and in humans with advanced AMD (Zinkernagel et al. 2017, Zysset-Burri et al. 2020) suggest that a distinct gut microbiome contributes to the development of late, neovascular AMD. No study has investigated the intestinal microbiome in patients with early AMD. Those studies in early AMD are of high importance as they potentially provide a gut microbiome signature reflecting stage of disease as well as a better understanding of the positive role of diet in disease pathomechanisms.

In humans, a healthy diet, rich in antioxidants such as a Mediterranean diet or supplementary antioxidants are the only known modifiable factors having a beneficial protective effect on progression of early AMD to advanced AMD (Merle et al. 2019, Raimundo et al. 2018, Age-Related Eye Disease Study Research Group 2001). Merle et al. (2019) demonstrated in a prospective cohort study consisting of two large European populations that the incidence of advanced AMD was modulated by a Mediterranean diet. Whether this result is due to effect of diet on microbial composition is unknown. The influence of light on the microbiome composition has been only investigated in rodents (Deaver et al. 2018). In mice, irregular ambient light exposure patterns alter gut bacterial composition and upregulate gene expression related to chronic inflammation (Deaver et al. 2018). This pathway is thought to disrupt the intestinal barrier thereby creating a pro-inflammatory environment (Deaver et al. 2018). Light signalling is also disrupted in early (Maynard et al. 2015) and late AMD (Maynard et al. 2017) due to photoreceptor damage affecting retinal photoreceptor classes and in particular, intrinsically photosensitive retinal ganglion cells (ipRGCs) that also receive light inputs from rod and cones for photoentrainment (Feigl and Zele 2014). Aberrant light signalling via dysfunctional ipRGC inputs may therefore contribute to gut microbiome dysbiosis and chronic inflammation in humans with AMD.

Published human studies have not controlled for or investigated the effect of diet or ambient light, two factors which are known modulators of the gut microbiome (Kolodziejczyk et al. 2019, David et al. 2014, Deaver et al. 2018, Godinho-Silva et al. 2019, Rowan et al. 2017) nor did they consider gut microbiome composition in early-stage AMD. Here, we investigate the gut microbiome in early AMD and whether a controlled, healthy diet affects its composition while controlling for ambient light and photoreceptor signalling.

## Materials and methods

### Study participants

All experiments were performed in accordance with the Queensland University of Technology (QUT) Human Research Ethics Approval (approval number: 1700001174) and the declaration of Helsinki; informed consent was obtained from all participants. Ten AMD patients (two females, eight males, age = 73.8 years ± 5.7) and ten age-similar healthy controls (four females, six males, age = 67.8 years ± 10.2) were recruited from the QUT Health clinics. All participants were in good health; systemic medication mainly included treatment of hypertension (four/ten) and one received treatment for gout. None of the participants were taking antibiotics at the time of the study. All participants were assessed for medical history and underwent a comprehensive ophthalmological examination including visual acuity assessment (Bailey-Lovie Log MAR chart), slit-lamp examination, tonometry (ic100; Icare Finland Oy, Finland), indirect ophthalmoscopy, central retinal and retinal nerve fiber layer (RNFL) thickness measurement with optical coherence tomography (OCT) (NIDEK RS-3000 OCT RetinaScan Advance; Nidek Co., Ltd., Japan) and fundus photography (Canon Non-Mydriatic Retinal Camera, Canon Inc., Japan) to exclude eye diseases other than early AMD, as well as to confirm the health status of the control participants (Table 1). Both groups had lenticular opacifications less than grade 2 (LOCS III) (Chylack et al. 1993), no anterior eye abnormalities and normal intraocular pressure (< 21 mmHg). In the AMD and control group, four (three in both eyes and one in the left eye) and two (in both eyes) participants respectively, had an intraocular lens (IOL) due to previous cataract surgery. AMD grading was performed as per AREDS classification (Table 1) (Age-Related Eye Disease Study Research Group 2001). There was no significant difference in central macular thickness (Mann-Whitney U = 171; *P* = 0.43) and RNFL thickness (independent t-test; t38 = 0.49; *P* = 0.63) between the groups (Table 1). None of the participants had diabetes, was obese, took proton-pump inhibitors or reported being gluten intolerant.

**Table 1.** Clinical characteristics of early AMD and control participants.

|  | Age | Gender | Visual Acuity<br>(Log MAR) | Lens<br>(LOCS) | OCT<br>macula | AREDS | Systemic<br>Morbidity |
| --- | --- | --- | --- | --- | --- | --- | --- |
| AMD 1 | 80 | M | 0.3 | IOL | 268 | 3D | RA |
| AMD 2 | 75 | M | 0.14 | 0 | 113 | 2C | Arrhythmia,<br>Urothelial<br>carcinoma |
| AMD 3 | 83 | M | 0.36 | IOL | 308 | 2C | Gout |
| AMD 4 | 81 | F | 0.14 | 1.0N | 263 | 1 | RA, HT |
| AMD 5 | 75 | M | 0 | IOL | 290 | 3C | Anxiety |
| AMD 6 | 71 | M | 0.1 | IOL | 274 | 1 | Osteoporosis,<br>HTN |
| AMD 7 | 77 | F | 0.3 | 1.5N | 257 | 2C | - |
| AMD 8 | 84 | M | 0.1 | 0.5N | 286 | 2B | - |
| AMD 9 | 77 | M | 0 | 0.5N | 272 | 2C | HTN |
| AMD 10 | 64 | M | 0.08 | 0 | 272 | 3B | HTN |
| **Mean** | **76.7** |  | **0.15** |  | **260.3** |  |  |
|  | Age | Gender | Visual Acuity<br>(Log MAR) | Lens<br>(LOCS) | OCT<br>macula | Systemic<br>Morbidity |  |
| Control 1 | 64 | M | 0 | 0.5N | 279 | - |  |
| Control 2 | 55 | F | 0 | 0 | 275 | HTN |  |
| Control 3 | 61 | F | 0 | 0 | 245 | - |  |
| Control 4 | 61 | M | 0.02 | 0 | 303 | - |  |
| Control 5 | 57 | M | 0 | 0 | 303 | - |  |
| Control 6 | 73 | F | 0 | 0.5N | 219 | HTN |  |
| Control 7 | 88 | M | 0 | IOL | 267 | Stroke,<br>Asthma |  |
| Control 8 | 80 | M | 0.1 | IOL | 279 | - |  |
| Control 9 | 71 | F | 0 | 0.5C | 244 | - |  |
| Control 10 | 71 | F | 0 | 0 | 233 | - |  |
| **Mean** | **68.1** |  | **0.01** |  | **264.7** |  |  |

LOCS: lens opacities classification system, IOL: intra-ocular lens, N: nuclear opalescence, C: cortical cataract, OCT: optical coherence tomography, AREDS: age-related eye disease study, RA: rheumatoid arthritis, HT: hypothyroidism, HTN: hypertension

### Diet

Diet can rapidly change microbiome composition in less than 5 days (David et al. 2014). All 20 participants received the same healthy diet over five days. The diet had a twofold purpose: 1) to normalize all participants to the same diet to reveal an inherent difference in the gut microbiome between controls and AMD participants that might be concealed due to the participants individual dietary regimes and 2) to unmask the mechanism through which a well-balanced healthy Mediterranean diet acts on the gut microbiome, given diet is proposed to be beneficial in preventing AMD progression (Merle et al. 2019). The diet consisted of a daily breakfast, lunch, dinner and snacks, delivered to the participant’s home and was based on a normocaloric Mediterranean diet (1500 kcal) prepared by a commercial kitchen (Lite n’ easy, Brisbane, Australia). All participants provided a fecal sample in the morning before starting the diet and on the next morning after completing the five-day diet. Drinks (e.g., water, coffee, tea, alcohol) were not provided and were recorded in a diary. Both groups did not differ in their consumption of liquids and had on average two-three coffees/tea/day (Independent t-test, *P* = 0.7) and one-two standard alcohol servings/day (Independent t-test, *P* = 0.8). One control participant had a probiotic drink once per /day and one AMD patient had a glass of milk every day.

### Determination of the diversity and the relative abundance of the gut microbiome

In total, 40 fecal samples (20 participants X 2 time-points, once before and once after diet) were collected to determine the diversity and the relative abundance of the gut microbiome. The fecal samples were stored at -20°C in the participant’s home freezer prior to extraction of microbial DNA (DNeasy PowerLyzer PowerSoil kit, Qiagen, Australia) according to the manufacturer protocols. The quantity and quality of the extracted DNA was determined using the Implen NanoPhotometer^®^ (LabGear, Australia). DNA samples were stored at -80°C until analysis. DNA was sequenced on the MiSeq 16S Metagenomics system using 300bp paired reads and the MiSeq v3 reagents which sequences the V3 and V4 regions of the 16S rRNA gene (Illumina, Australia). Post-sequencing, Illumina’s BaseSpace metagenomics workflow was used to perform taxonomic classification based on the Greengenes database. This workflow demultiplexes indexed reads, generates FASTQ files, and then classifies reads at several taxonomic levels ranging from kingdom to species. A Calypso compatible .csv file containing the microbial operational taxonomic units (OTUs) was uploaded to the online Calypso program for data analysis and visualization (http://cgenome.net/calypso).

### Actigraphy

To measure ambient light exposure (lux) and activity, all participants wore a wrist actigraphy (GENEActiv Original, Activinsights, UK) continuously for five days during the dietary intervention. The data were sampled in 1-minute epochs after removing the values during non-wear times(Joyce et al. 2019). The individual daily day and night light exposure was calculated using the sunrise/sunset time in Brisbane, Australia (http://www.ga.gov.au/) and averaged across the five days. The data were corrected for the underestimations of true luminance by GENEActiv using the toolbox provided by our laboratory (Joyce et al. 2019).

### Photoreceptor signalling

To confirm aberrant light signalling in our participants cohort with AMD, we quantified the progressive photoreceptor loss (Curcio et al. 1996) in AMD, including melanopsin expressing ipRGCs (Maynard et al. 2015, Maynard et al. 2017),(Hattar et al. 2003). The afferent pupil response provides an objective, non-invasive and *in vivo* means to directly quantify the function of outer and inner retinal photoreceptors in response to light (Gamlin et al. 2007, Markwell et al. 2010, Kelbsch et al. 2019). We determined melanopsin photoreceptor function using pupillometry and the post-illumination pupil response (PIPR) as per established methods in our laboratory (Zele et al. 2011, Feigl and Zele 2014, Adhikari et al. 2015, Kelbsch et al. 2019). In short, following 10 minutes adaptation to the darkened room (< 1 lux), we measured the consensual pupil response from the right eye while stimulating the left eye in Maxwellian view using a custom-designed pupillometer (50° diameter stimulus; 20.1 mm retinal image diameter; 15.5 log.quanta.cm^-2^.s^-1^ corneal irradiance) (Feigl and Zele 2014, Adhikari et al. 2015). We used two stimuli with high melanopsin excitation; a 1 s short wavelength (blue) stimulus (λ_max_ = 460 nm; full width at half maximum [FWHM] = 23 nm; melanopsin excitation = 7997.57 melanopic lux(Lucas et al. 2014) and a medium wavelength (green) stimulus (λ_max_ = 519 nm; FWHM = 34 nm; 6985.78 melanopic lux), the latter being less attenuated by ocular media, and a third stimulus with low melanopsin excitation as a control (long wavelength (red); λ_max_ = 630 nm; FWHM = 16 nm; 4.05 melanopic lux). The effect of circadian variation was minimized by performing pupillometry between 10am and 5pm(Zele et al. 2011). Measurements were repeated twice for each wavelength at both visits (12 measurements/participant and 240 measurements in total). The PIPR amplitude was defined as the pupil diameter at 6s after stimulus offset relative to the baseline pupil diameter (Kelbsch et al. 2019).

## Statistical analysis

Primary outcomes were microbiome diversity and abundance before and after diet. Secondary outcome measures included the melanopsin-mediated PIPR to quantify ipRGC function and environmental light exposure (actigraphy).

Microbial diversity and abundance analysis for all samples was done using Calypso V8.84 (http://cgenome.net/calypso) (Zakrzewski et al. 2017). Calypso enables quantitative visualisations, statistical testing and multivariate analysis of the metagenomics sequence data. Total sum normalization (TSS) was used for data normalization which normalizes data by dividing feature read counts by the total number of reads in each sample. The method converts raw feature counts to relative abundance. The data were further transformed by TSS combined with square root transformation (Hellinger transformation). After data filtering and normalisation, various statistical tests, such as one-way Analysis of Variance (ANOVA) with post hoc tests (unpaired t-tests), and sample clustering methods such as Canonical Correspondence Analysis (CCA) multivariate were used. The core microbiome differences between sample groups (AMD vs controls, before and after diet) were determined using a > 0.40 relation for the top 300 most abundant taxa. Microbial alpha diversity (comparison within sample) was calculated using Shannon’s index, evenness, richness, Simpson’s index and Chao 1 tests. Between sample (beta) diversity was tested using CCA (samples clustered by location) and Anosim (tests if sample community profiles are significantly different between sample groups). In addition, Calypso’s Group functionality was used to compare abundance of bacterial Phyla and Species across sample groups (ANOVA parametric statistical testing) with significantly different Phyla and Species shown using bar charts with *P* < 0.05.

GraphPad Prism (GraphPad Software, Inc., CA, USA) was used for statistical analysis of pupil and actigraphy data. Frequency distributions were evaluated with the D’Agostino and Pearson omnibus normality test. The PIPR amplitudes before and after diet were compared with a paired t-test (normal data) or the Wilcoxon test (non-normal data) (95% confidence interval, *P* < 0.05). To compare the pupil and light exposure data between the participants with AMD and healthy controls, an independent t-test (normal data) or the Mann-Whitney test (non-normal data) was applied.

## Results

### Faecal sample analysis

Metagenomic sequencing of 40 faecal samples was done to detect differences in the baseline microbiome between the two groups, and after normalization for diet. Bacterial diversity was determined at genus and phylum levels. We found no significant differences in alpha and beta diversity at all levels between and within the AMD and control groups before and after diet (Figure 1A-C). Figure 2 shows the taxonomic characterisation of bacterial phyla present in both groups before (A) and after diet (B) which was not significantly different between the groups before and after diet. To determine which bacterial species were significantly differentially abundant in AMD patients, analysis of the 300 most abundant OTUs before and after diet was performed. There were significantly differentially abundant bacterial species before and after diet between AMD and control groups (Figure 2C, D). This analysis also revealed that independent of diet, three bacterial species, namely, *Clostridium termitidis, Bacteroides gallinarum* and *Bacteroides finegoldii* were significantly differentially abundant in early AMD patients compared to the control group (Figure 2C, D). Interestingly, *Sutterella* sp. that is associated with inflammatory intestinal disease (Mangin et al. 2004, Gophna et al. 2006) was only significantly abundant in AMD compared to controls at baseline (Figure 2C) but not after the diet. Species recognized as commensals/beneficial bacteria in the gastrointestinal tract (Wexler 2007, Wang et al. 2019) such as *Parabacteroides distasonis* and *Bacteroides nordii* were significantly more abundant in the AMD group after diet compared to controls (Figure 3D, F) but not before diet (Figure 3C, E).

**Figure 1.**
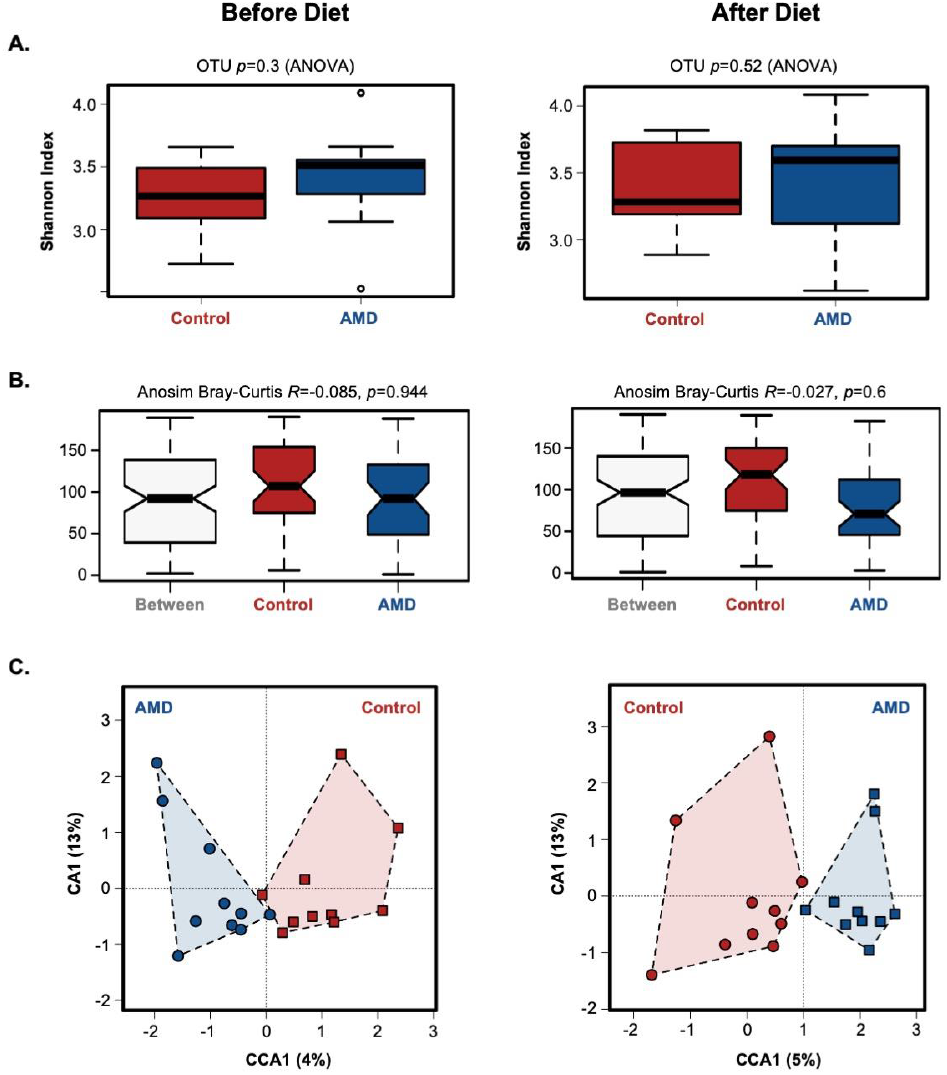
A-C. Diversity of bacterial species at OTU level in AMD compared to control participants before (MD1, C1) and after diet (MD2, C2). **A**. Shannon index with 25^th^ and 75^th^ percentiles and median indicating species diversity (alpha diversity), **B**. Beta-diversity (diversity between groups) and **C**. canonical correspondence analysis (CCA) for analyses of clustering of samples do not show significant differences.

**Figure 2.**
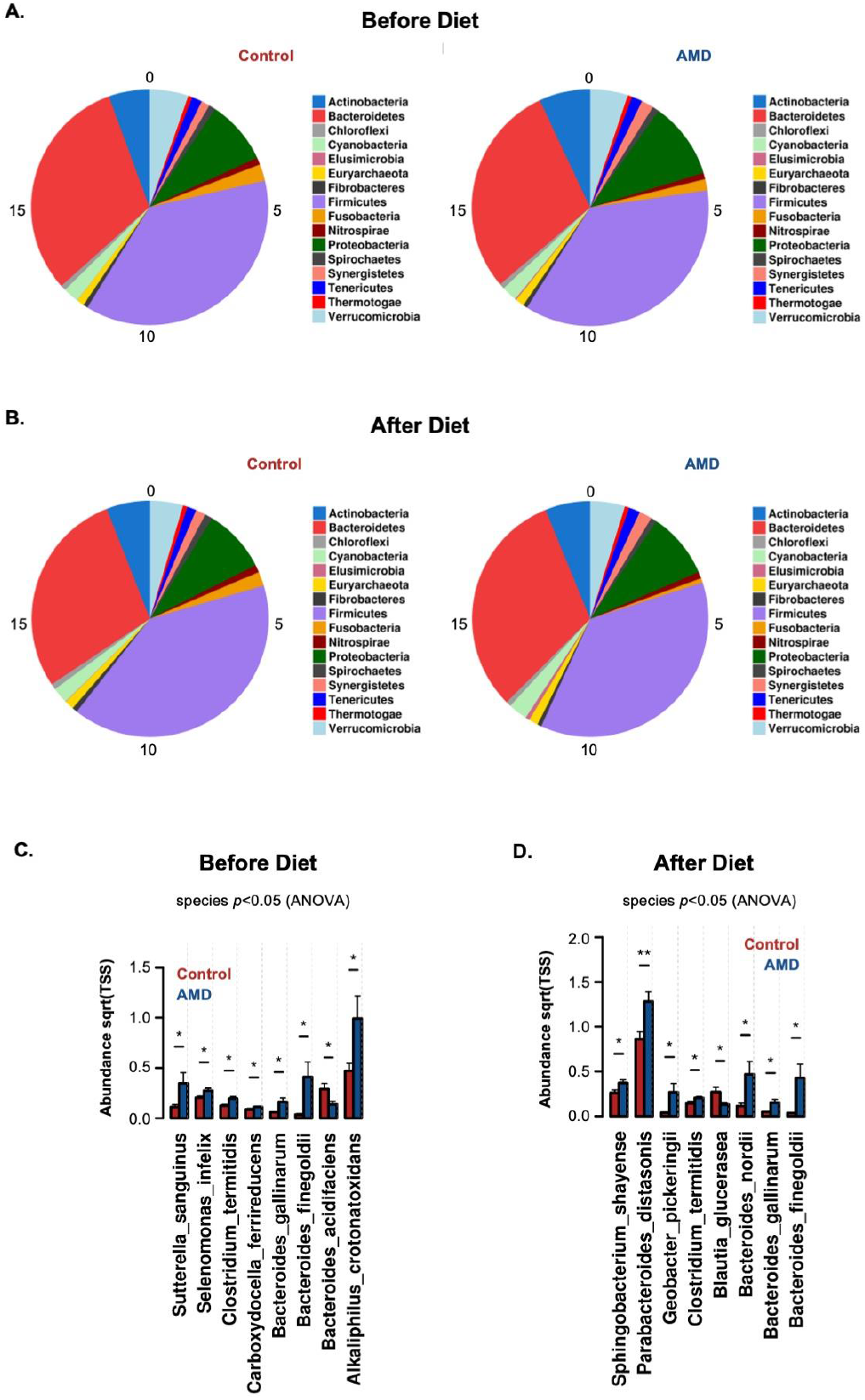
A-D. Taxonomic characterisation (Phylum) in AMD and control groups **A**. before and **B**. after diet. Core analysis of the 300 most abundant OTUs between AMD and controls **C**. before and **D**. after diet demonstrating significantly abundant pro-inflammatory and protective species, respectively in early AMD patients (*p<0.05).

**Figure 3.**
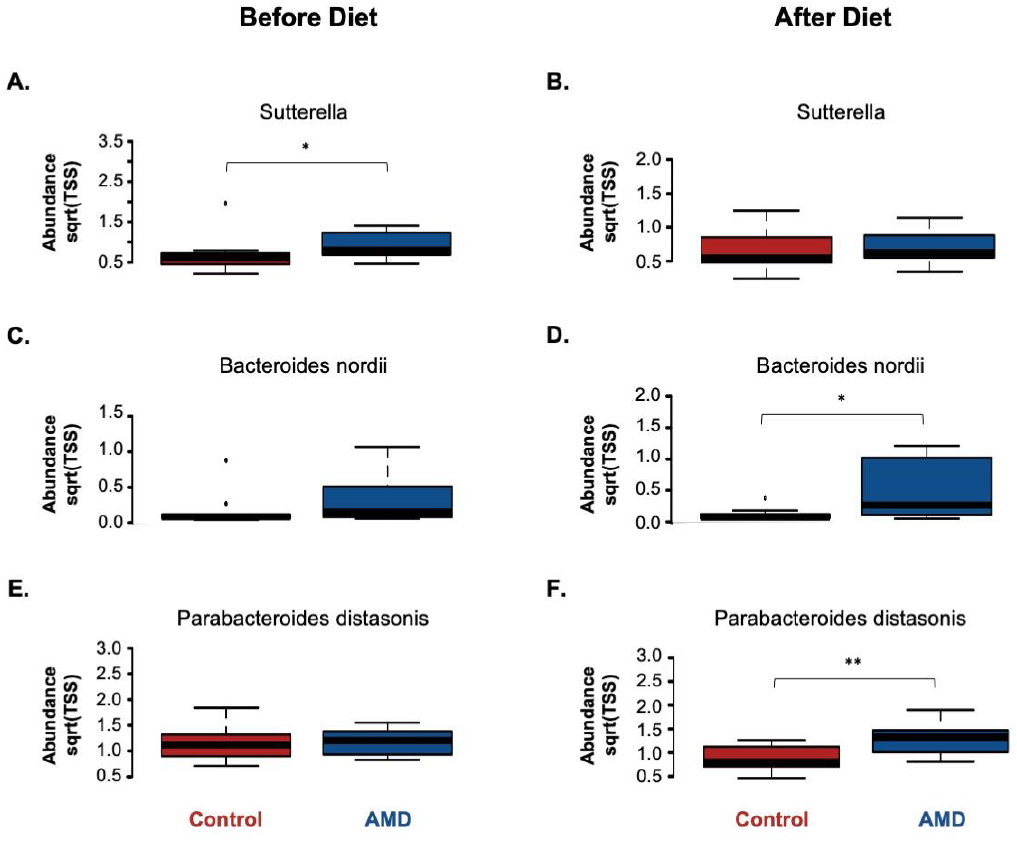
A-F. A. Box plots representing the mean abundance (+/-SD) of *Sutterella* sp. associated with AMD before diet (*Kruskal-Wallis p<0.5); blue are AMD patient samples and red are control samples . **B**. Box plots representing the mean abundance (+/-SD) of *Sutterella* sp. showing no differential abundance in AMD after diet. **C**. Box plots representing the mean abundance (+/-SD) of *Bacteroides nordii* showing no differential abundance in AMD before diet. **D**. Box plots representing the mean abundance (+/-SD) of *Bacteroides nordii* showing significant abundance in AMD after diet (*Kruskal-Wallis p<0.5). **E**. Box plots representing the mean abundance (+/-SD) of *Parabacteroides distasonis* showing no differential abundance in AMD before diet. **F**. Box plots representing the mean abundance (+/-SD) of *Parabacteroides distasonis* showing significant abundance in AMD after diet (*Kruskal-Wallis p<0.05); blue are AMD patient samples and red are control samples.

### Melanopsin function and ambient light exposure (actigraphy)

The PIPR was significantly reduced in early AMD compared to healthy controls (Figure 4B, C; Independent t-test; Blue light: t_18_ = 5.3; *P* < 0.0001; Green light: t_18_ = 3.1; *P* = 0.01). Actigraphy analysis demonstrated that AMD and control groups had similar environmental light exposure (Mann-Whitney U = 1020592; *P* = 0.47) and activity (comparison in sedentary, light, moderate, and vigorous activity between the groups using independent t-tests: t_18_ = 0.30 to 1.51; *P* = 0.15 to 0.77) during the dietary intervention (Figure 4A), therefore excluding any effect of differences in ambient lighting that could affect the microbiome in AMD patients.

**Figure 4.**
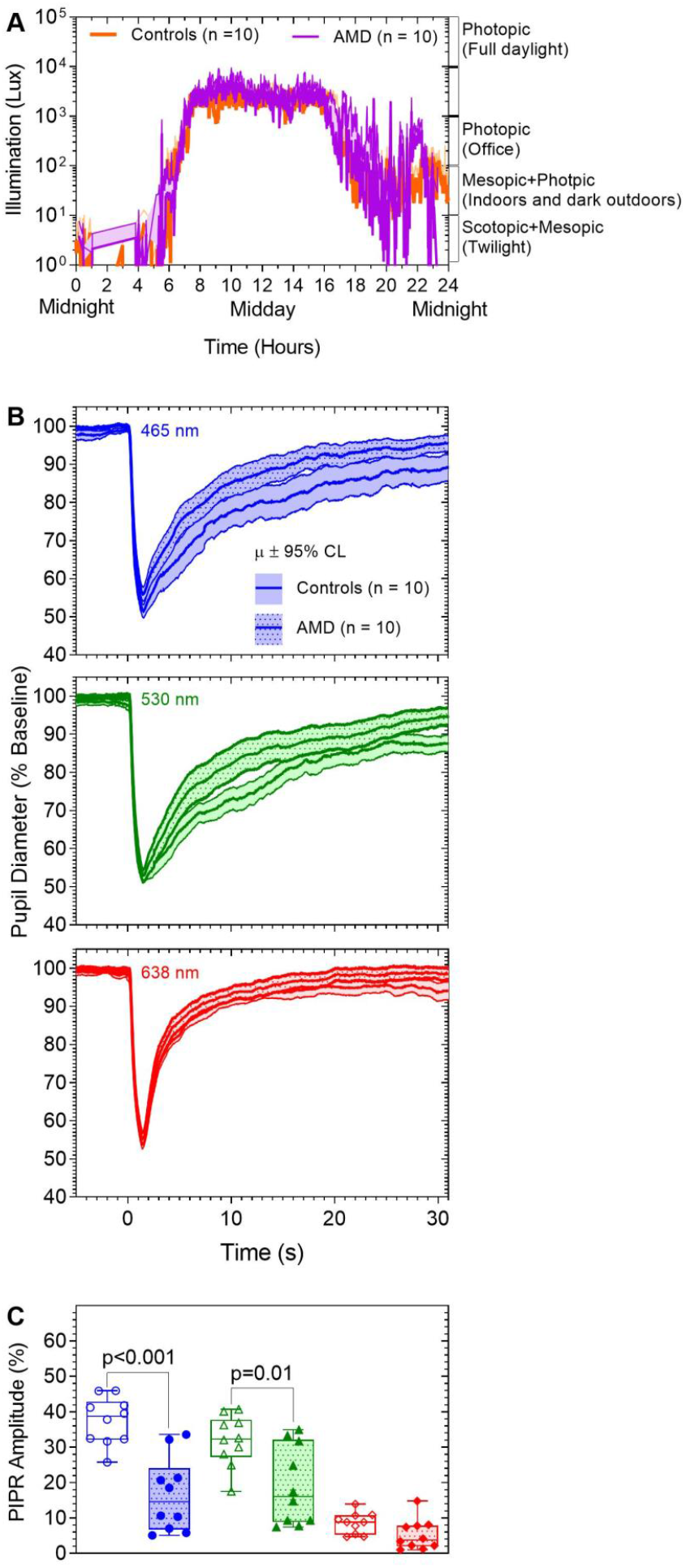
A-C. A. The environmental light exposure as measured with actigraphy over the period of the diet shows no significant difference between the AMD and the control group. **B**. The average and 95% confidence limit of the pupil light reflex (PLR) in response to a high (blue) melanopsin excitation (465 nm), a lower (green) melanopsin (530 nm) stimulus, and a red (638 nm) stimulus with no melanopsin excitation demonstrates a reduced PIPR response (pupil returns to baseline) for the blue and green melanopsin stimuli in AMD (dotted) compared to the controls (filled). There is no significant difference for the red stimulus between the groups. **C**. Box and whisker plots shows the median interquartile and range of the 6sPIPR amplitudes for red, blue and green wavelength stimuli for in early AMD (filled symbols) in comparison to controls (unfilled symbols). The 6s PIPR amplitudes for blue and green stimuli were significantly reduced in AMD compared to the control group.

## Discussion

We provide the initial evidence of a distinct gut microbiome in early AMD. We demonstrate a significantly differential abundance of the pro-inflammatory species S*utterella* before controlling for diet (Figs 2C, 3A). Independent of diet, the species *Clostridium termitidis, Bacteroides gallinarum* and *Bacteroides finegoldii* were significantly differentially more abundant in the early AMD compared to the control group (Figs 2C, D). After normalizing for diet, we found significantly differentially more abundant commensal bacteria (*Parabacteroides distasonis* and *Bacteroides nordii)* in AMD (Fig 3D, F).

We show that both groups had the same ambient light exposure (Figure 4A). The difference between AMD and control participants, however, was the presence of ipRGC dysfunction in early AMD (Figure 4B, C) that can lead to aberrant photoreceptor signalling of the ambient illumination to the central circadian clock in the SCN. Whether this light disruption may contribute to bacterial dysbiosis which has only been shown in animal studies (Deaver et al. 2018) needs to be determined in humans.

While the microbial composition affecting the bacterial abundance has been investigated in patients with advanced AMD (Zinkernagel et al. 2017, Zysset-Burri et al. 2020), these studies did not control nutritional intake or quantify light exposure. Previous studies in advanced AMD (Zinkernaglel et al 2017, Zysset-Burri et al. 2020) found a relative abundance of *Anaerotruncus, Oscillibacter, Ruminococcus torques, Negativicutes* and *Eubacterium ventriosum*. The differences in microbiome composition in our study in early AMD compared to existing studies in late AMD may also reflect the different stages of AMD. S*utterella* sp. are associated with chronic inflammatory bowel disease (Mangin et al. 2004, Gophna et al. 2006). AMD is also considered as chronic inflammation (for review (Feigl 2009)) and further studies are required to investigate the nature of intestinal damage caused by *Sutterella* sp. in early AMD patients. After the healthy diet, commensal bacteria abundance increased in AMD. This suggests a positive effect of diet in facilitating beneficial species dominating the intestinal microbiome that could play a protective role in AMD.

We observed that the beneficial bacteria were significantly differentially more abundant despite defective ipRGC signalling after the diet, suggesting that diet could potentially override the effect of circadian disruption on the gut microbiome, at least in early disease. Follow on studies will be required to evaluate if supplementary light exposure that boosts the defective melanopsin photoreceptor response has additional beneficial effects, as does diet, to modulate the microbiome towards an anti-inflammatory environment.

We do not believe age or comorbidities impacted on our data as the age difference between both groups was only marginally significant (t_14.3_ = 2.3, p = 0.041) and none of our participants had an uncontrolled chronic condition, were obese or had diabetes (Table 1). We also controlled for dietary as well as light exposure which are all main modulators of the microbial composition in the gut. Here, we have normalised both AMD and control groups to the same diet to demonstrate a “temporary” effect on microbial composition. We acknowledge that such effect must be tested over a longer period to demonstrate whether a healthy diet can directly affect retinal pathology to show no progression of drusen to advanced, wet or dry AMD.

## Conclusion

In summary, we provide the initial evidence of a distinct gut microbiome in a controlled, well-defined group of people with and without early AMD. We demonstrate that a healthy diet can modify the relative abundance of the gut microbiome towards more beneficial species over a short period of time. We infer that a potential mechanism of diet on AMD progression may be through changes in the gut microbial composition.

## Data Availability Statement

The data presented in this study can be found in online repositories. The name of the repository and accession number will be provided in the final document.

## Ethics Statement

All experiments were performed in accordance with the Queensland University of Technology (QUT) Human Research Ethics Approval (approval number: 1700001174) and the declaration of Helsinki; informed consent was obtained from all participants.

## Author contributions

BF, AJZ and FH conceived and designed the study and wrote the manuscript. Text editing by all co-authors. BF was the lead in all experiments and was supported by AJZ and PA (vision data) and FH (microbiota data). Under the guidance of BF, AJZ and FH, PA performed pupil data and actigraphy collection analysis. Under guidance of BF and FH, NJ performed microbiota analysis.

## Funding

Supported by the Australian Research Council Discovery Projects ARC-DP170100274 (BF, AJZ) and an Australian Research Council Future Fellowship ARC-FT180100458 (AJZ).

## Conflict of Interest

The authors have no competing conflict of interest.

## Notes

### Competing Interest Statement

The authors have declared no competing interest.

